# Dynamic clustering of genomics cohorts beyond race, ethnicity—and ancestry

**DOI:** 10.1101/2023.08.04.552035

**Authors:** Hussein Mohsen, Kim Blenman, Prashant S. Emani, Quaid Morris, Jian Carrot-Zhang, Lajos Pusztai

## Abstract

**Background:** Recent decades have witnessed a steady decrease in the use of race categories in genomic studies. While studies that still include race categories vary in goal and type, these categories already build on a history during which racial color lines have been enforced and adjusted in the service of social and political systems of power and disenfranchisement. For early modern biological classification systems, data collection was also considerably arbitrary and limited. Fixed, discrete classifications have limited the study of human biodiversity and disrupted widely spread genetic and phenotypic continuums across geographic scales. Relatedly, the use of broad and predefined classification schemes—e.g. continent-based—across traits can risk missing important trait-specific genomic signals.

**Results:** To address these issues, we introduce a dynamic approach to clustering human genomics cohorts on a trait-specific level and without using a set of predefined categories. We tested the approach on whole-exome sequencing datasets in ten cancer types and partitioned them based on germline variants in cancer-relevant genes that could confer cancer type-specific disease predisposition. Results demonstrate clustering patterns that transcend discrete continent-based categories across cancer types. Functional analysis based on cancer type-specific clusterings also captures the fundamental biological processes underlying cancer, differentiates between dynamic clusters on a functional level, and identifies novel potential drivers overlooked by a continent-based clustering model.

**Conclusions:** Through a trait-based lens, the dynamic clustering approach reveals genomic patterns that transcend predefined classification categories. We propose that coupled with diverse data collection, new clustering approaches have the potential to draw a more complete portrait of genomic variation and to address, in parallel, technical and social aspects of studying human biodiversity.

## Introduction

### 1.1. Historical perspective

The earliest scientific attempt to use race as a category to classify human beings dates back to the seventeenth century. In a 1684 essay titled, “A New Division of the Earth, According to the Different Species or Races of Men Who Inhabit it,” French physician Francois Bernier categorized human beings into five types, the last of which, the Sámi people, he described as “little stunted creatures” and “wretched animals” [1]. Swedish botanist Carl Linnaeus, dubbed as the founder of modern taxonomy, published decades later (1735) the first edition of *Systema Naturae*, in which he created a system with four categories instead, and based his categories on geography and skin color. In the tenth edition (1758), he expanded the system and confounded physical with personality and social traits based on his interpretation of the humoral theory that links geography and climate to skin color and good and bad character [2]. In this work, Linnaeus loaded his classifications with prejudice and crafted a hierarchy placing *Homo sapiens europaeus*, a category he color-coded as white, on top, while using descriptions such as “harsh face,” “careless,” “stubborn,” “lazy,” “greedy,” and “ruled by caprice” to describe *Homo sapiens americanus*, *afer* and *asiaticus*, color-coded respectively as red, black, and yellow [1, 3, 4]. Linnaeus also added a separate category he called *Homo sapiens monstrosus*, in which he mostly included humans with various birth defects and mythical “humans” such as giants from Patagonia [2–5].

Bernier and Linnaeus suggested different sets of categories and imbued their human classification systems with imagined hierarchical value judgement. Their imposition of a few, fixed, distinct and discrete categories reduced the complexity of human biodiversity and shaped subsequent classification systems. In early systems of classification, descriptions often depended on the philosophical (and political) choices of the classifiers, technological limitations, and economic factors including trade routes [2, 6]. For instance, the routes involving Sweden and the Netherlands during Linnaeus’ time shaped his considerably arbitrary choice to describe peoples of specific geographies but not others. Further, Linnaeus relied on anecdotal and written accounts of his students and of missionaries, mercantilists, travelers, and slave traders, and did not travel himself outside of western Europe [2, 4].

Early biological classification systems were also venues for the use of emerging modern science to demarcate human difference in the service of power during times of colonial expansions and the Atlantic slave trade. Institutions and individuals exerted intentional efforts to create racial classification systems in modern science, which opened the door for racialized hypothesis generation. In a stark yet far from lone example, the Bordeaux Royal Academy of Science announced an essay contest in 1739 to study “the degeneration of Black hair and Black skin.” The announcement was made a year after the regional assembly of Bordeaux endorsed the existence of enslaved Black people on French soil, and as recently described in “Who’s Black and Why: A Hidden Chapter from the Eighteenth-Century Invention of Race,” before members of the Bordeaux Academy decided to invest Academy prize money in the company that ran the French slave trade in the African continent, Compagnie Perpetuelle des Indes [7]. Another example of a (re)defined color line to benefit the interests of chattel slavery before its abolition in the United States is the introduction of “one drop laws,” according to which a person was categorized Black if they had a known “trace of Black blood” in their ancestry. As a result, in the words of anthropologist Nina Jablonski, skin color was “no longer the necessary and sufficient criterion for race classification” [2] (for more examples, see [8, 9] and references listed in Box 1-1 of [10]).

Changing meanings of race categories continued to reflect and drive political and social transformations. Since the beginning of the first U.S. census in 1790, for example, racial groupings in the census have changed more than twenty times [1]. Notably, race categories and their meanings also vary across national borders [1, 10]. Biology and medicine are susceptible to societal and cultural influences, and scientists are engaged in a bi-directional process of being influenced by social and cultural concepts that co-shape interpretations of nature, and scientific interpretations that in turn influence social order [11, 12]. Scientific attempts to formulate biological classification systems of race continued in the nineteenth and twentieth centuries and were muddled, again, with confusion. In the words of anthropologist Fay-Cooper Cole during the opening of the “Conference on Racial Differences,” which was held in 1928 at the National Academy in Washington, D.C. and attended by opponents and proponents of eugenics, the term “race” was “frequently used in three or four ways in the same article,” and there existed “a great deal of confusion in the use of the word.” Eugenics, which propagated race science for more than six decades, declined during the 1930s and 1940s after it faced strong scrutiny and criticism within scientific circles and in response to Nazi eugenical horrors. Yet, the use of racial classification systems to study human biodiversity continued [13]. The understanding of human biodiversity has progressed since then, however, and further highlighted their unreliability.

### 1.2. Further limitations

The reduction of human biodiversity into a small set of skin color-coded race categories implies the existence of distinct skin color lines. However, skin color is a continuum influenced by climate, genetics, and the intensity and seasonality of ultraviolet radiation [2]. Similar skin colors can also result from convergent adaptation in response to similar selective pressures, rather than from genetic relatedness [2, 14], and further analysis and data collection demonstrated the prevalence of continuous rather than discrete skin color distributions (e.g. [15, 16]). Further, recent comprehensive efforts centered on diverse genomics data have demonstrated (i) the complexity and prevalence of continuums on the genome level (e.g. [17]), and (ii) the sharp limitations of broadly predefined classification—be it at the level of socio-political categories such as race and ethnicity, or broad geographic ones such as continental ancestry (e.g. [18]).

While genetic ancestry is a concept that describes a partial relationship of a person with their genealogical history, it can still be subject to significant limitations as a classification criterion. First, there is no single criterion to define ancestry, and categories can take geographic (e.g. South Asian or Central American), geopolitical (e.g. Zambian or Italian), or cultural (e.g. Brahmin or Lemba) meanings [14]. Second, geographic ancestry categories might be muddled with imprecise conflation with race categories, and their descriptors can be distortive of time and space (e.g. references to Asian ancestry might exclude the entirety of or wide regions within West, Central, East or South Asia, and nationality-based categories might refer to ancestors during the period that preceded the very formation of respective countries). Third, continent-based categories reduce the high levels of genomic complexity within each continent and might inadvertently imply purity within continents when communicating results. Fourth, separate categories impose discreteness on continuums between continents [14, 17, 19–21] that have long been connected by land (e.g. Asia and Europe, or today’s Asia and Africa through the Sinai Peninsula), in technology, or both.

Further, single category assignments to individuals ignore the multitudes of personal belonging. While many individuals choose to affiliate with multiple groups for personal or cultural reasons [14], it is also highly common for individuals to have a “mixed genetic ancestry” with respect to sets of predefined categories (e.g. 97.3% of individuals are associated with a median of four ancestry categories in [18], in consistence with [22]). Importantly, this “mixed ancestry” can be observed on the individual level and does not have to reflect population stratification. Further, significant amounts of genetic ancestry labeled as “Western Asian,” for example, is present in samples with origins ranging from present-day Morocco to Mongolia, and from England to Ethiopia, that is, in Asia, Europe and Africa [18].

### 1.3. Dynamic, trait-specific germline clustering

Given the limitations of predefined classification systems, and recognizing the wide range of phenotypes and the complexity of genomic variation, we propose a dynamic approach that generates trait-specific clusterings of genomics cohorts. The approach builds on an earlier idea from an exchange between biological anthropologist Frank B. Livingstone and evolutionary biologist Theodosius Dobzhansky on generating clusterings on the gene(s) level (see [23] and Chapter 9 of [13]), and expands it in light of the wide advances in genomic data collection, measurement of genomic variation, and interpretation of the genomic basis of complex traits. The approach is also motivated by multiple factors. First, the genomic basis of different traits is encoded in different loci of the human genome, and the loci relevant for a single trait, ranging from one to many in number, cover only a small portion of the whole genome. Second, biological and physiological processes are shared among individuals within and across any number of groups— however ancestrally defined. Third, especially when common germline polymorphisms are involved in part or in full in trait predisposition, the genomic variants are significantly shared across continental regions. Relatedly, a single nucleotide polymorphism (SNP) can be concurrently classified “rare” in multiple regions, and the distribution of classifications depends on available data [24].

Fourth, evolutionary forces might be acting on trait-specific genomic regions in parallel in distant geographies. Fifth, as predefined ancestral labels concurrently bear geopolitical, historical, and social meanings, their assignment to categories used to study genomic biodiversity, and particularly predisposition to disease, opens the door for prolonging a history of stigmatizing entire communities [25].

Finally, the disruption of observed continental clines overlooks inter-continental patterns related to a trait of interest. This raises a core question on the goal of clustering cohorts: if two individuals in distant geographies have similar genomic markers and phenotypic expression corresponding to a trait, e.g. both are right-handed, should they be in the same cluster when studying the genomic basis of handedness, or separate ones? Should “populations” be determined based on a gene(s) (or trait) of interest, or the whole genome? What are the limitations of clustering based on the whole genome, in a fragmented and data-scarce continental setting, when a trait is affected by only a small subset of genomic regions? Further, it is also quantitatively well-established that selected features or clustering criteria strongly affect resulting “populations” (i.e. clusters or clines), and consequently reshape the starting point from which to discover—or miss—patterns and generate hypotheses [26, 27].

The dynamic clustering approach takes a different angle to classifying biodiversity by grouping individuals based on predisposition to a trait under study—herein a cancer type. An individual’s membership to a cluster depends on their genomic sequence at a specific set of regions known to be associated with the trait. Number of clusters, which are *de facto* neutrally labelled, is determined according to the dataset and specific trait under study.

In cancer, germline (inherited) predisposition is mediated by deleterious mutations in several dozen high penetrance cancer-relevant genes and probably a combination of individually low penetrance variants. Different genes are associated with different degrees of risk, and with variable cancer-specificity [28]. Further, germline alterations require additional acquired (somatic) mutations for malignant transformation [29–31]. We hypothesize that clustering cancers based on their germline variants in cancer type-specific loci transcends predefined continental categories. We expect clusterings to vary across cancers—in terms of number of clusters and sample-cluster membership—to reflect human biodiversity and heterogeneity across complex traits. We also note that this dynamic (i.e. trait-specific) approach moves beyond the notion of local ancestry at a single locus as it can simultaneously consider any set of coding or non-coding regions associated with a trait, and it can scale to accommodate newly acquired knowledge on the genomic basis of the trait under study.

## Results

### 2.1. Overview

To study predisposition to ten of the most common cancer types [32], we utilized germline data from the Cancer Genome Atlas (TCGA) [28, 30] and ancestral category values from the The Cancer Genetic Ancestry Atlas (TCGAA) [33]—which are based on comparisons with the 1000 Genomes [34], HGDP [35] and HapMap [36] projects. In studied cohorts, TCGAA uses categories that refer to continental and sub-continental regions. For clarity, we hereafter refer to them collectively as continent-based.

We first generated the trait-specific clusterings of each cohort, using the TCGA terminology for cancer types, BRCA, COAD, KIRC, LIHC, LUAD, LUSC, OV, PAAD, PRAD, and READ, using all nonsynonymous SNPs within different sets of COSMIC genes known to have germline association with each cancer type (Methods, Supplementary Table 3). Next, we focused on the SNP subset predicted to have high functional impact within each cancer type’s samples as a basis for dynamically generating clusters. We then assessed the performance of three algorithmic clustering approaches (K-means, DBSCAN, and HClust) to identify generated clusters in multidimensional scaling (MDS) plots. Finally, we identified potentially overlooked somatic driver genes in each TCGA cohort based on dynamic clustering in comparison with drivers identified using the continent-based model, and performed a functional genomic analysis to assess their biological and clinical importance in the context of cancer.

### 2.2. Beyond continent-based categories

Upon dynamically clustering based on trait-specific regions—herein cancer type-specific germline COSMIC genes, a number of visual patterns emerge. First, the number of clusters varies per cancer type (1-8 clusters), strongly reflecting known genomic heterogeneity in cancer [28, 30, 37] (Figure 1). Second, and despite the relative lack of diversity in TCGA datasets, clusters transcend continent-based categories in all cancer types to include samples with “African,” “East Asian,” “European,” and “Other” ancestral labels within clusters. Third, this pattern is also observed in colon and rectum cancers, known to be associated with high disparities in incidence and outcome [38, 39], with one and two clusters (Figure 1b and 1j), respectively.

**Figure 1.**
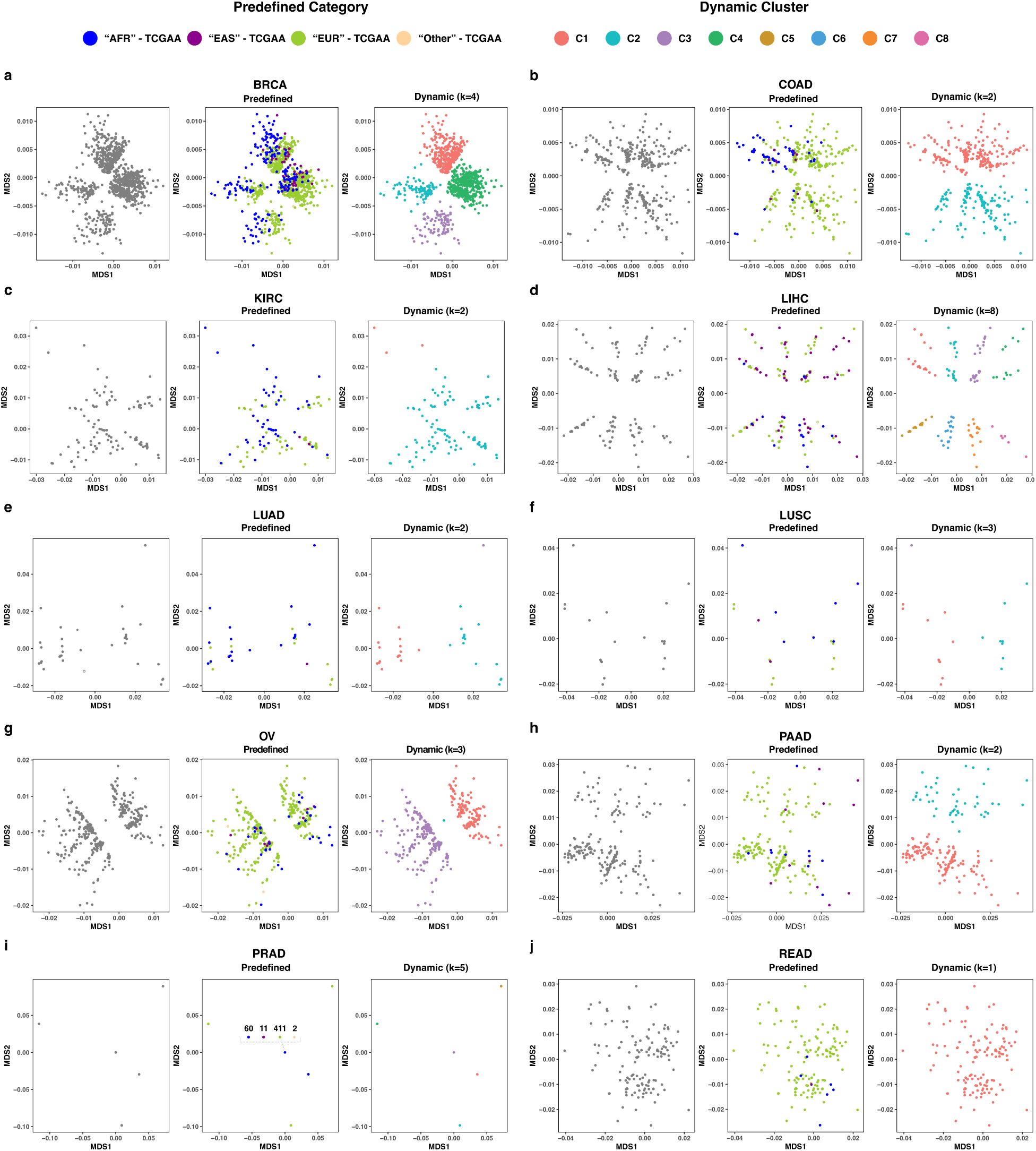
Multidimensional scaling (MDS) plots of dynamically generated clusters for ten TCGA cancer cohorts. Cancer type-specific dynamic clusters transcend predefined continent-based categories. Dynamic cluster numbers (i.e. C1, C2, … C8) correspond to disjoint sample subsets within each cancer cohort.

### 2.3. High functional impact compact clusters

Next, we selected subsets of SNP variants with high functional impact on protein function (HFI) within the COSMIC genes corresponding to each TCGA cohort as a basis for clustering (see Methods). This selection led to SNP subsets with n = 1 (PAAD) to 269 (BRCA). HFI-based dynamic clusters transcend continent-based boundaries and exhibit two notable patterns. First, the number of HFI-based clusters is higher, on average, than clusters based on all nonsynonymous SNPs of COSMIC genes (e.g. PAAD with two COSMIC-based vs four HFI-based clusters in Figure 1h and Figure 2a, respectively; HFI-based clusters for other cancer types in Supplementary Figure 1). Second, subsections within select cancer types tend to be compact and often include samples with highly similar or identical HFI variant patterns due to the smaller number of loci used for clustering in these cohorts. Dots corresponding to distinct samples overlay each other, resulting in single dots each representing a subcluster (e.g. LUSC-HFI in Figure 2a and KIRC-HFI in Supplementary Figure 1).

**Figure 2.**
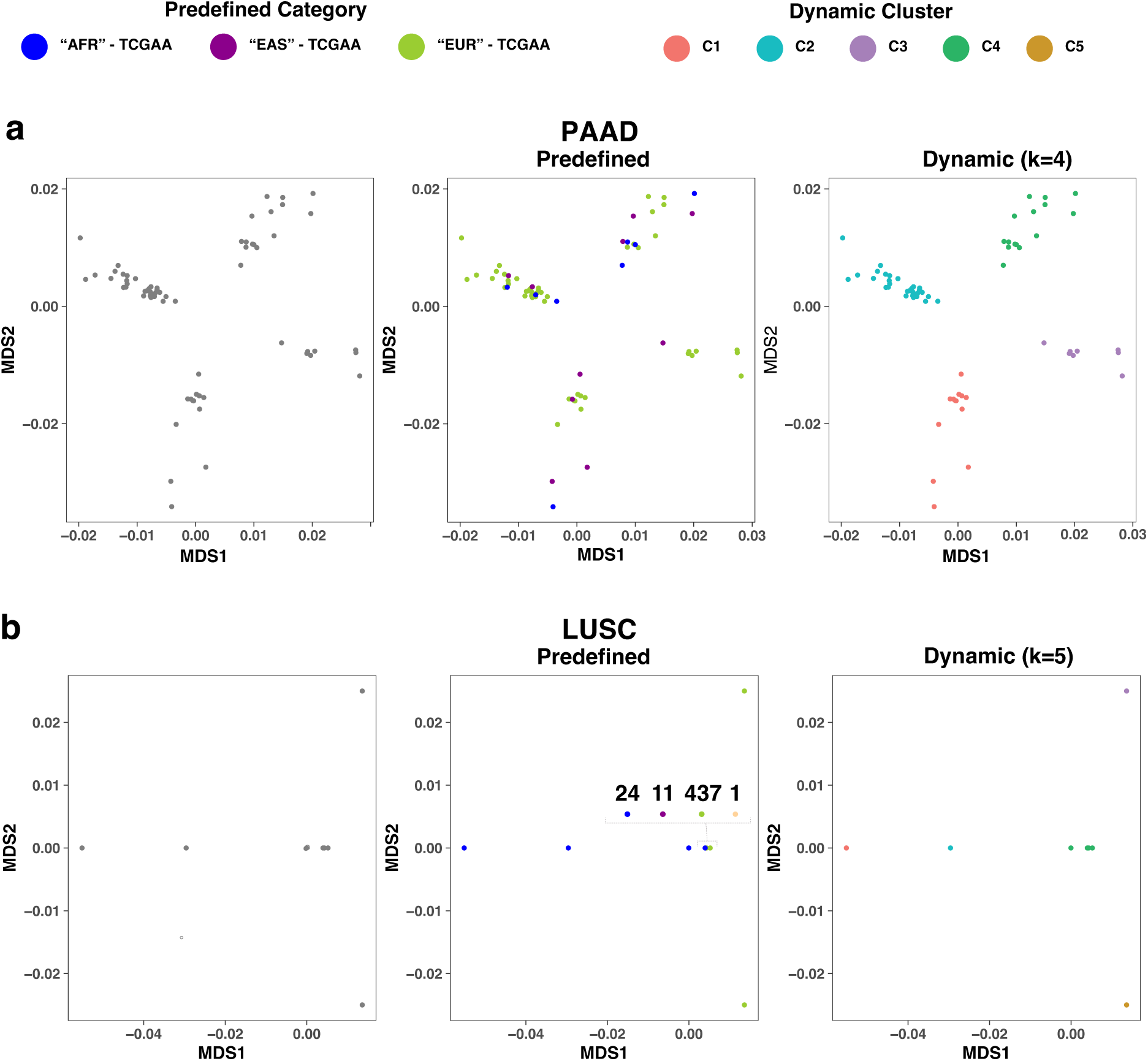
MDS plots of dynamically generated clusters based on high functional impact germline variant subsets. (a) PAAD results demonstrate a higher number of clusters in HFI-based results compared to ones based on all nonsynonymous variants in the COSMIC-based setting in Figure 1h. (b) LUSC results demonstrate compact clusters with a high number of samples demonstrating similar or identical variation patterns in HFI subsets.

### 2.4. Human aid improves algorithmic clustering

Algorithmic approaches can generate different clusterings of the same dataset, and their performance primarily depends on data distribution and cluster definition [27, 40]. To algorithmically identify clusters, we ran the K-means algorithm with a predefined k = 4 (number of selected TCGAA continent-based categories) and an “optimal” k chosen based on visual inspection of cancer type-specific plots. Given the varying number of clusters across cancer types, predefined-k clustering performed poorly across multiple cancer types (Figure 3a-d). Similarly, dynamic-k K-means faced challenges in accurately identifying clusters across cohorts (e.g. LIHC-COSMIC, Figure 3e).

**Figure 3.**
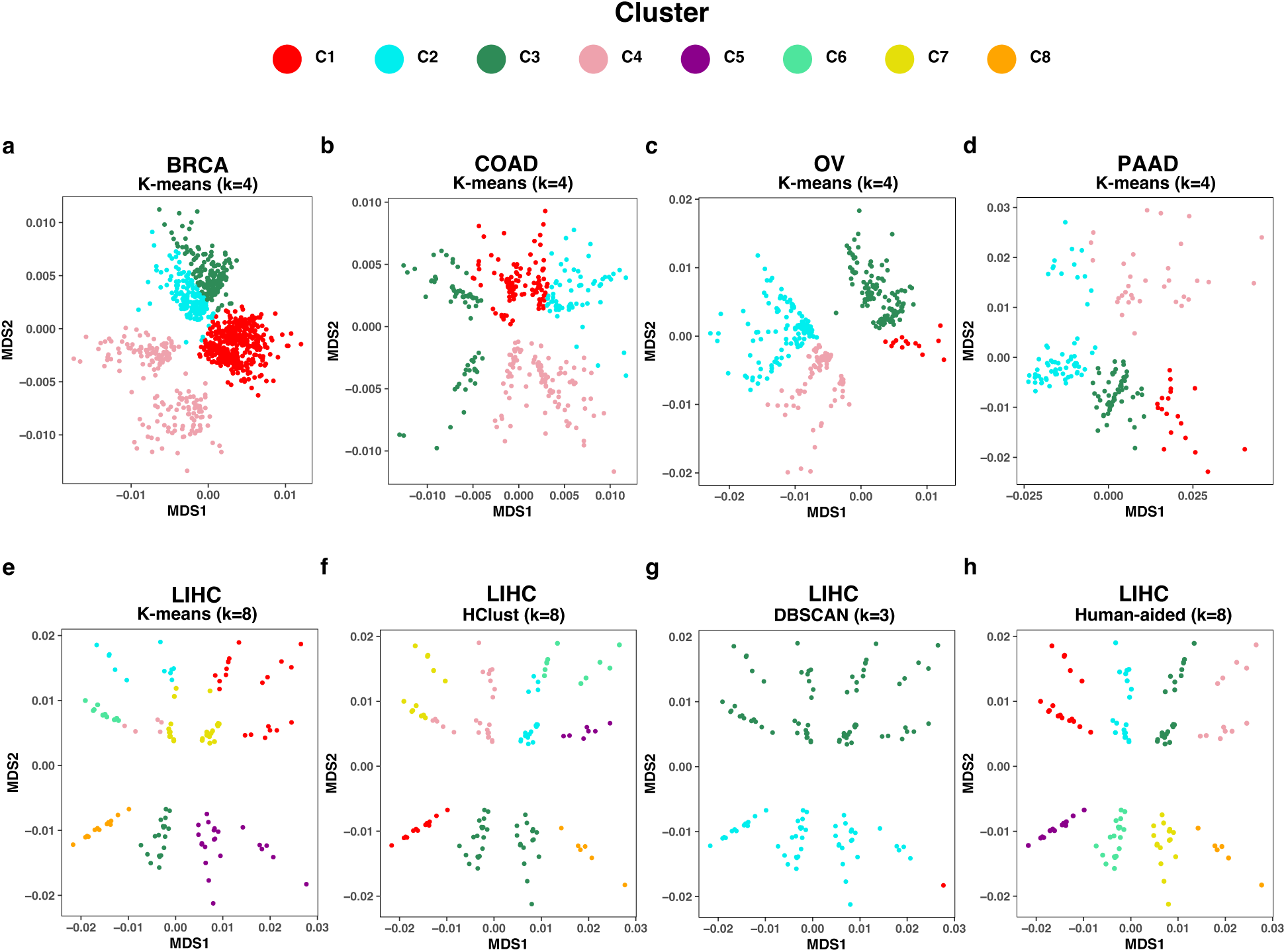
Algorithmic and human-aided identification of dynamic clusters. K-means results with predefined-k = 4 fails to identify COSMIC-based clusters in (a) BRCA, (b) COAD, (c) OV, and (d) PAAD among other cancer types. Dynamic-k results also demonstrate the failure of (e) K-means, (f) HClust, and (g) DBSCAN to identify COSMIC-based clusters in LIHC, highlighting the need for (h) human-aid in cluster identification.

Similarly, clustering results from two other algorithms, DBSCAN (Density-based Spatial Clustering of Applications with Noise) and HClust, performed poorly in multiple cohorts. HClust, which stands for agglomerative hierarchical clustering—herein used with complete-linkage, is a bottom-up approach that starts with individual points as separate clusters and iteratively merges most similar clusters until a predefined number of clusters is met (e.g. k = 8 clusters for LIHC-COSMIC in Figure 3f). Like K-means, HClust correctly identified only a subset (i.e. two) of the eight clusters in LIHC-COSMIC. DBSCAN, which is known to identify dense clusters and outliers in low-density regions, partitioned LIHC-COSMIC results into three clusters (Figure 3g), a number decided algorithmically based on input parameters (Methods): two large clusters, each roughly with four of the dynamic clusters, and a third cluster that includes distant samples the algorithm considered outliers (in red).

In sum, and while specific algorithms perform better than others at identifying clusters, algorithmic clustering does not seem to suffice for both COSMIC- and HFI-based settings. As a result, human intervention to attain more precise results remains central (e.g. Figure 3h). We also note that in certain instances, multiple “optimal” numbers of clusters in the same plot can exist, and the choice remains centered on experimental goals and the problem under study (e.g. the four dynamic clusters of BRCA-COSMIC in Figure 1a being alternatively considered two larger diagonally-separated clusters in the same plot).

### 2.5. Dynamically-identified cancer drivers

We next explored the biological significance of clusters identified by the dynamic clustering approach. Particularly, we focused on identifying potential cancer type-specific somatic driver genes. We used MutSigCV to identify potential somatic drivers based on whole exome sequence data corresponding to each dynamic cluster generated on germline variant sets (COSMIC genes and HFI). COSMIC-based clusters yielded 98 potential drivers across cancer types, and HFI-based clusters 109 drivers. Both lists included tens of known drivers from a more comprehensive list by Bailey *et al.* which relied on multiple computational and experimental tools [41]. These include *KRAS*, *TP53*, *PIK3CA*, *BRCA1*, *PTEN*, *CDH1*, *RB1*, *PTEN*, *FOXA1*, *SPOP*, and *VHL* (for full lists, see Supplementary Tables 1 and 2). COSMIC- and HFI-based lists also include 23 and 36 cancer type-specific novel potential drivers, respectively, which are overlooked if analysis is performed on continent-based clusters (Figure 4a). Among these genes are known drivers listed in [41], including *APC*, *CBFB*, *B2M*, *CDKN2A*, and *RPL5*.

**Figure 4.**
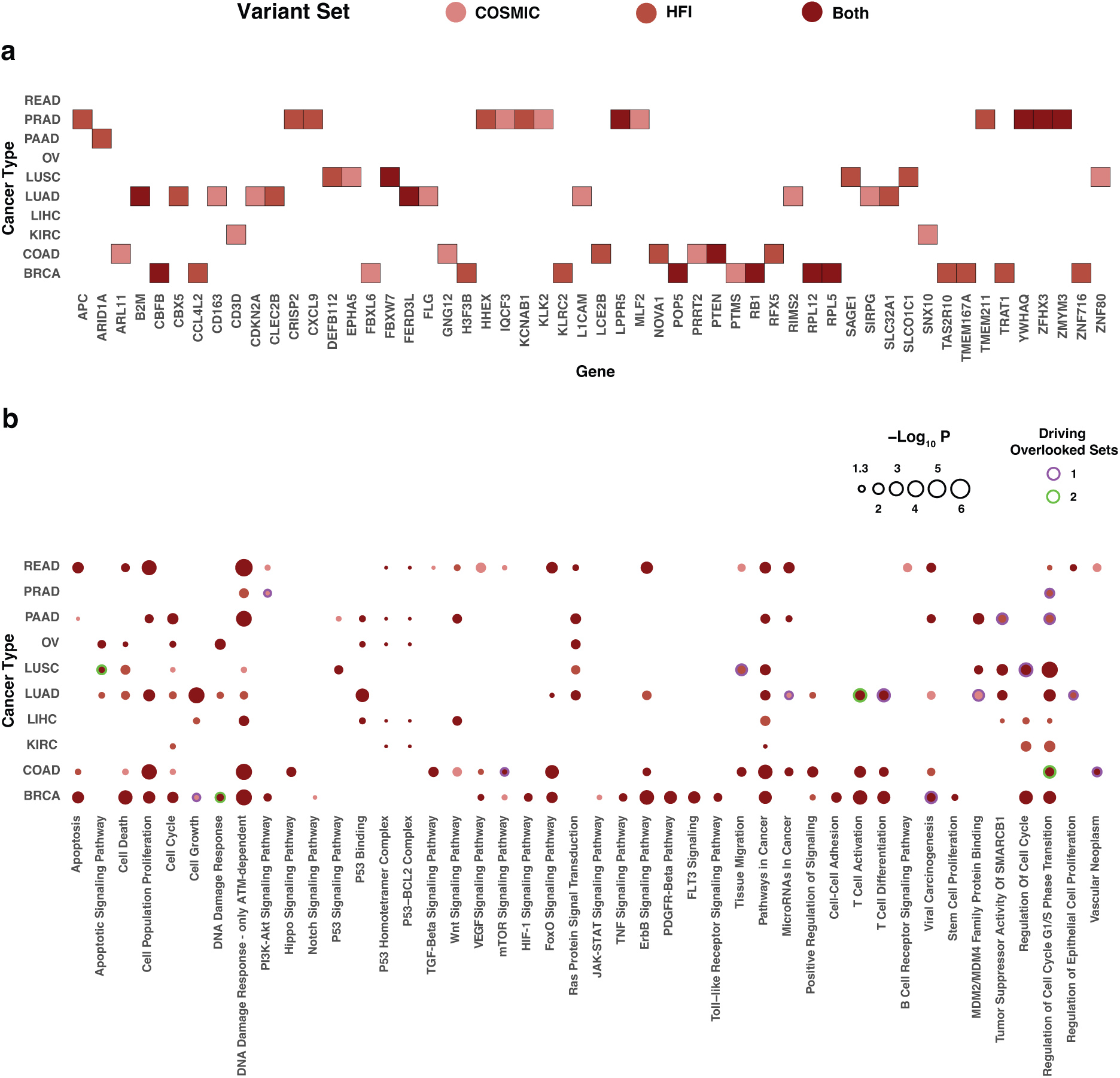
Known and potential driver genes identified based on dynamic clustering. (a) Dynamic cluster-based genes overlooked by the continent-based model. Each of the listed genes was identified in at least one COSMIC- or HFI-based dynamic cluster and none of the clusters based on predefined continent-based categories. (b) Dynamic cluster-based drivers associate widely with known cancer pathways. Genes overlooked by the continent-based model drive a subset of these associations in one (purple border) or both (green) settings centered on the COSMIC- and HFI-based variant sets.

We then investigated the functional importance of COSMIC- and HFI-based driver gene lists. Given their significant coverage of known drivers, enrichment analysis of both full lists point to known oncogenic pathways, including the majority of signaling pathways listed in Sanchez-Vega *et al.* [42]. These pathways include cell cycle alongside Hippo, Notch, PI3K/Akt, TP53, TGFB and WNT signaling. Among other important pathways and processes are apoptosis, HIF-1, mTOR, TNF, JAK-STAT, VEGF, and FoxO signaling (Figure 4b), and pathways named after several cancer types and other diseases and infections (e.g. Epstein-Barr and Kaposi sarcoma-associated herpesvirus virus infections). We compared enrichment results in the resulting dynamically-generated lists with and without novel drivers overlooked by the continent-based model. The inclusion of novel genes allows dynamic clusters to refer to biological processes, pathways, and entities with strong effect on cancer etiology (purple and green borders) such as apoptotic signaling in LUSC, DNA damage response in BRCA, mTOR signaling in COAD, and multiple terms pertaining to cell adhesion, tissue migration, cell cycle, and cell proliferation across multiple cancer types.

### 2.5. Dynamic clusters are distinguishable by clinical and functional cancer signifiers

Dynamic clusters vary by the composition of their underlying germline variants. To investigate the biomedical significance of the resulting clusterings, we tested for the associations between each of the clusters, compared to its all its counterparts within a cancer cohort, with clinical variables and gene expression patterns in TCGA. In the LUAD-COSMIC setting, the age at the first pathologic diagnosis is lower in cluster 1 (C1) than 2 (mean = 64.4 and 66.6 years, respectively; Wilcoxon rank-sum test, *p_adj_* < 0.05; Figure 5a). In LIHC, C1 of the COSMIC setting shows a significant enrichment for high tumor grade samples, and C1 of the HFI setting for late tumor stage ones (Figure 5b and 5c, respectively; Wilcoxon rank-sum test, *p_adj_* < 0.05), with similar results that vary among clusters in LIHC and LUSC cohorts (Supplementary Figure 2).

**Figure 5.**
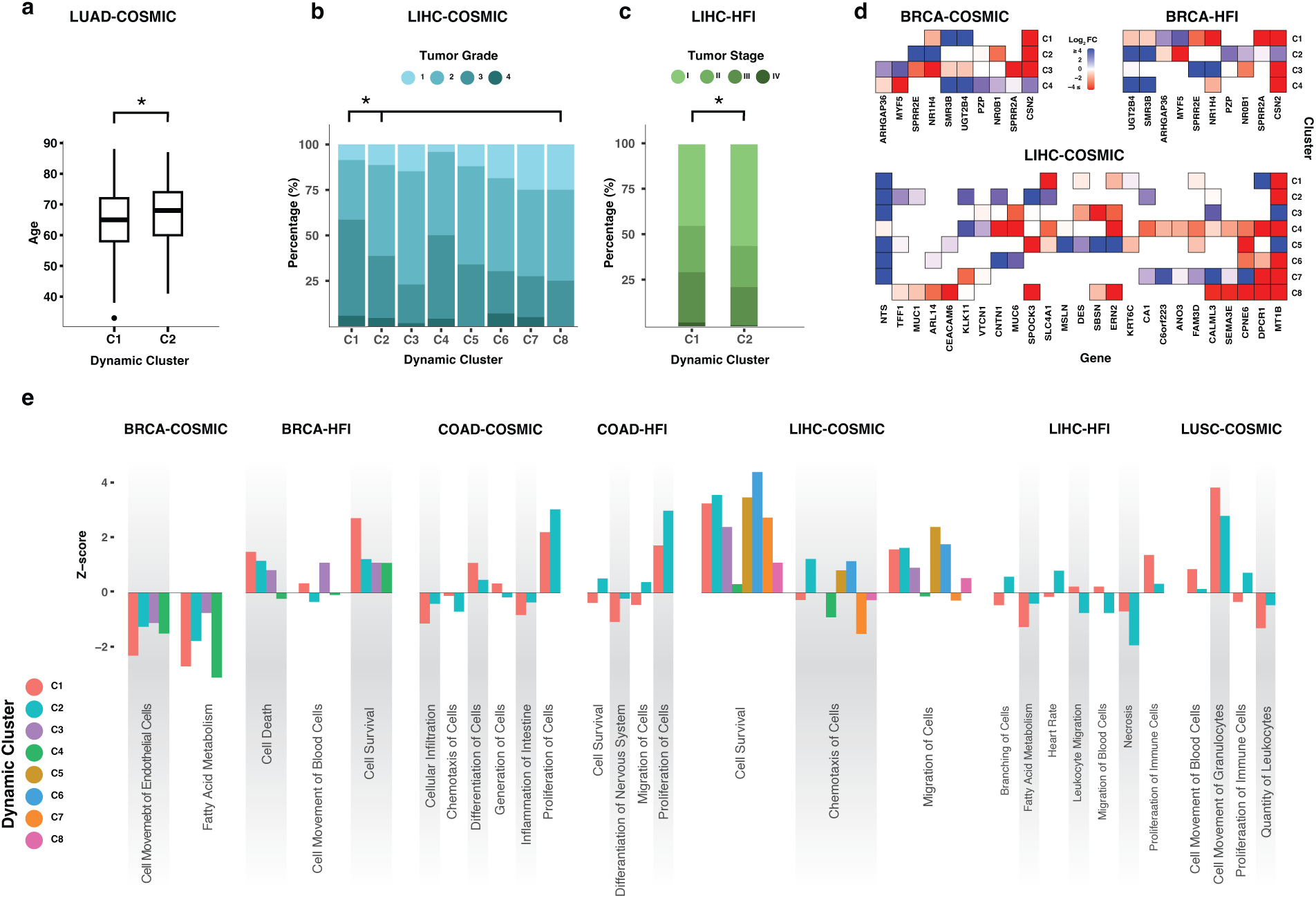
Dynamic clusters across cancer types highlight clinical and functional associations. a) Dynamic cluster 1 (C1) based on the COSMIC subset in LUAD (LUAD-COSMIC) shows statistically significant lower age cancer onset than that of the second cluster (C2; *p_adj_* < 0.05) b) C1 in LIHC-COSMIC includes samples with higher tumor grade compared to other clusters combined (*p_adj_* < 0.05). Clusters C5 and C8 include no Grade 4 samples. c) C1 shows more advanced tumor stage samples in LIHC-HFI compared to C2 (*p_adj_* < 0.05) d) Genes significantly expressed (FDR < 0.1) in opposite directions among clusters of BRCA-COSMIC, BRCA-HFI and LIHC-COSMIC highlight potential functional roles associated with different clusters e) Gene programs with known association to cancer are collectively expressed in different magnitudes and directions between dynamic clusters across cancer types and settings.

We then shifted attention to analyzing differential gene expression patterns across cancer types and settings. At the individual gene level, opposite expression levels are detected among clusters of the same cohort (Figure 5d). Notably, such patterns correspond to genes with reported associations with tumorigenesis, patient survival, metastasis, and cell proliferation. These include *CSN2* [43], *NROB1* [44], and *NR1H4* [45] in both COSMIC and HFI settings in BRCA, and *MT1B* [46], *MUC1* [47], and *KLK11* [48] in LIHC-COSMIC. At the gene program level, we used Moonlight [49] to identify programs that are collectively differentially expressed in each cluster compared to matched “normal” samples. Within different cohorts, the magnitude and direction of expression mark significant differences among clusters. These include, among others (Figure 5e; Supplementary Figure 3), a negative expression (Z-score < 0) of cell death in only one out of four clusters in BRCA-HFI (C1), of migration of cells in only two out of eight in LIHC-COSMIC (C4 and C7), and of cell survival and migration of cells in one of the two clusters in COAD-HFI (C1); a significantly higher expression of proliferation of cells in one cluster in each of COAD-COSMIC and COAD-HFI (C2 in each); and a significantly lower expression of cell survival in LIHC-COSMIC (C4), quantity of leukocytes in LUSC-COSMIC (C2), and necrosis and fatty acid metabolism in LIHC-HFI (C1 and C2, respectively).

### 2.6. Dynamic clusters are distinguishable by non-cancer signifiers

In addition to cancer-focused gene programs, we investigated the biological significance of specific genes significantly expressed in only one dynamic cluster within each setting (|Log_2_ FC| > 2). Biological enrichment analysis of these gene sets revealed essential and non-cancer-focused biological processes that distinguish different clusters. These include, among others (Figure 6, Supplementary Figure 4), multiple processes related to neuronal response to stimulus in cluster 2 (C2) of COAD-COSMIC, non-cancerous diseases and infections in C1 of LUSC-COSMIC, and neuron development in C1 of LIHC-HFI. Other processes that closely pertain to cancer from an essential point of view include ones revolving around immune response in C2 of READ-HFI, and cell differentiation, which recurrently emerged from individual clusters within multiple cancer types and settings (i.e. label “Across Clusters” in Figure 6).

**Figure 6.**
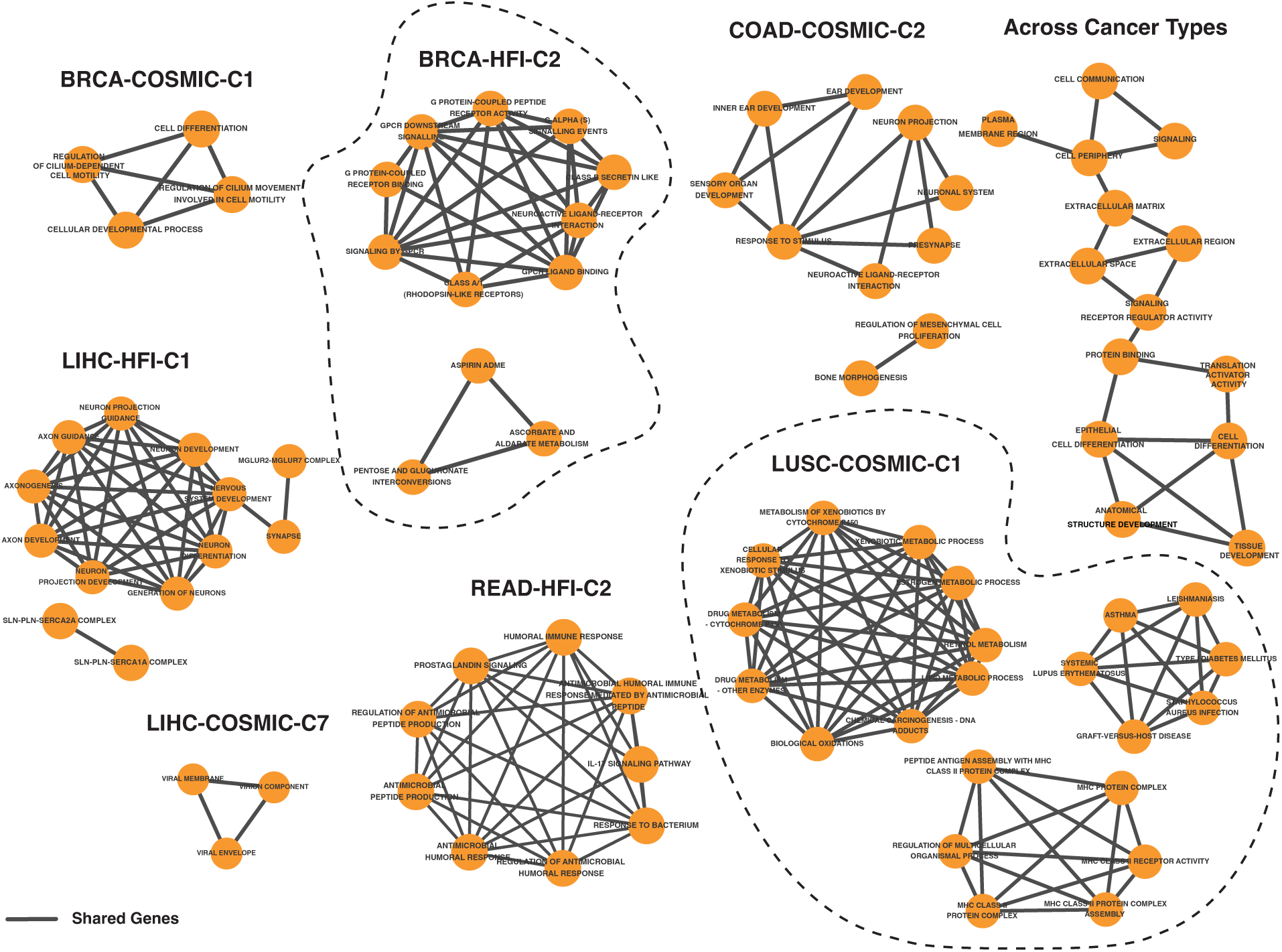
Genes expressed in single clusters within each cancer type and setting highlight related biological and clinical associations beyond cancer (e.g. asthma and lupus in LUSC-COSMIC-C1 and neural development in LIHC-HFI-C1). Resulting associations share a subset of their underlying genes (i.e. edges), and a number of biological processes recurrently emerges across cancer types and settings (i.e. “Across Cancer Types,” top-right).

## Discussion

Coupled with recent wide genomic data availability, existing, complex, and ever-changing human biodiversity engenders the need for multiple perspectives to studying genomic traits. We introduce a dynamic clustering approach that centers, in parallel, technical and social aspects of studying human biodiversity by utilizing a lens that focuses on trait-based genetic similarity. This approach does not lock samples within geography-based ancestry categories or average genomic patterns across the whole genome regardless of the trait under study. It also recognizes continuums and clusters when they exist at the trait-specific level (e.g. eight clusters for COSMIC-based LIHC vs one continuous cluster for READ in Figure 1). When we apply the dynamic approach to germline data of the TCGA cancer cohorts, emerging clusters transcend predefined continent-based categories. When we examine the somatic mutational patterns in the resulting dynamic germline-based clusterings, we identify tens of known and potential cancer driver genes, many of which were overlooked by the continent-based model. Results based on trait-specific clusters also capture the fundamental biology associated with the hallmarks of cancer and associate clusters with clinical and biological signifiers [50].

The dynamic approach has broader implications for how biodiversity is observed and classified. The use of racial classification systems in science has long been contested given their history that is rife with confusion, technological limitations, and enforcement in service of colonialism. Race, itself, is an idea rather than a discovery; an idea invented to impose systems of control and discrimination that continue to shape today’s social realities [51]. Enforced color lines disrupted continuous clines of variation and often shifted to serve political goals rather than to describe patterns of biodiversity. Further, race categories are usually collected to comply with civil rights reporting guidelines or for social and administrative purposes, but the racial categorization systems were not designed for genetic studies [2, 14]. Similarly, ethnicity categories can be centered on culture, social norms, religious beliefs, or language rather than genetic ancestry, and their use in genetic studies can lead to inaccurately reported results. Ethnicity categories are malleable concepts that can change in different times or circumstances irrespective of hereditary lines [14, 18].

A sharp decreasing trend in using race categories in genomic studies has been reported [52]. While genetic ancestry categories resemble, in their biological aspect, a direct reflection of a partial inheritance of genetic material across generational lines, they can also suffer from limitations in semantics, in space, and in time. Ancestry categories can be based on geographic, geopolitical, social, or cultural elements, and the process of imposing clear lines among predefined populations is rife with social and technical limitations. These limitations are heightened when categories are quite broad (e.g. continent-based), when individuals identify with more than one category, and when the labels might further open the door for enforced stigmatic associations of disease on entire communities [25]. As a result, it is generally advisable to cluster genomics cohorts only when justified by the research question rather than by default [10, 19], and to place social implications of the research at the heart of the design process rather than as an afterthought [53].

Given the compound nature of human biodiversity (i.e. continuums and clusters), a broader sampling of human genetic diversity—with the careful selection of categories during the data collection phase, if and as needed or obligated—remains highly central to more clearly understand the patterns of genetic variation (see Chapter 5 of [10] and [54]). In fact, it is through this type of diverse data collection efforts that continuous patterns of variation have been elucidated on a wider scale [17, 20]. Relatedly, alleles that increase the susceptibility to a disease can be present across multiple geographies. The notion of dynamic clustering can be carried over to genome-wide analyses as well. For example, in the case of methods that study one variant at a time—such as QTL identification and GWAS—it is possible to limit the SNP-based clustering to some neighborhood of each variant to consider how the local genetic relatedness of individuals can be accounted for in enrichment analysis.

While certain genomic patterns might be identified in a given dataset based on a given predefined model, this type of models is not necessarily the only route towards this type of identification. Equally importantly, as we demonstrate, other patterns can be missed when relying on broad categories or when considering complex traits with loci distributed across the genome. Broad stratification, whether driven by discriminatory legacies embedded in genetic practice or normalized experimental design and data collection, might obfuscate genomic patterns that transcend predefined categorical boundaries. Diverse datasets and new and existing quantitative approaches that can transcend predefined categories—by utilizing or being able to incorporate trait-specific regions, clines, estimated relatedness matrices (e.g. [55, 56]), principal components (e.g. [57–60]), or other chosen means—are hence crucial to approach different types of genomic studies (e.g. ones for gene discovery or other types of genomic studies described in [10]), to detect various patterns of genomic variation, to address theoretical and applied challenges (e.g. gene-environment interactions, pleiotropy, false positive control), and to leverage data generated using different technologies (e.g. deep sequencing and GWAS). These efforts have the potential to draw a more complete portrait of the genomic bases of traits all the while navigating the entangled relationships between science and society [11, 61–63].

## Methods

### Genomic Datasets

We used TCGA germline data from the BRCA (n=1072 samples after data processing), COAD (445), KIRC (514), LIHC (360), LUAD(513), LUSC(503), OV (556), PAAD (182), PRAD (488), and READ (164) studies. We used the MC3 somatic dataset [64] for functional enrichment analysis and the ancestral labels from the TCGAA Project (http://fcgportal.org/TCGAA/) [33] filtered according to the recommendations in [65].

### SNP Selection

For COSMIC-based SNP sets, we selected autosomal SNPs annotated as nonsynonymous, stop-gain, and stop-loss ClinVar database annotations [66]. For HFI subsets, the functional impact of missense germline variants within a cancer type’s exome samples was determined using MetaSVM [67] and annotations by ClinVar, when available. We considered a missense variant high-functional impact if classified as Deleterious by MetaSVM or listed as Pathogenic/Likely-Pathogenic in ClinVar. We used MetaSVM scores from the dbNSFP database which includes pre-calculated function impact scores for 75,931,005 human non-synonymous single-nucleotide variants [68]. Loss-of-function (LoF) variants, including frameshift indels, stop gain, and stop loss variants, were also considered high-functional impact as well as variants annotated as high-confidence LoF in gnomAD [69] or Pathogenic/Likely-Pathogenic in ClinVar. We only included autosomal variants that met quality control measures described in the source paper by Huang *et al.* [30]. The number of selected SNPs in each COSMIC and HFI variant set is available in Supplementary Table 4.

### Driver Gene Identification

Potential driver gene identification was performed on the MutSigCV [70] v1.3.4 server available at https://www.genepattern.org/modules/docs/MutSigCV. Genes with FDR < 0.1 were deemed statistically significant potential drivers.

### Clinical Variable Analysis

Clinical TCGA data was obtained from [71]. Continuous and ordinal variable comparisons (i.e. for age, tumor grade, and tumor stage) were performed using the Wilcoxon rank-sum test in a one-vs-all configuration on clusters with > 5% of samples within a cohort and COSMIC or HFI setting, with Bonferroni correction and *p_adj_* < 0.05 significance level.

### Gene Expression Analysis

Gene expression data corrected for batch effect and study-specific bias were downloaded from RNAseqDB [72] at https://github.com/mskcc/RNAseqDB. Differential expression analysis at the gene program level Moonlight v1.20 [49], and at the gene level was performed using edgeR v3.36 [73] (*q* < 0.05, |Log_2_ FC| > 2).

### Enrichment Analysis

Enrichment analysis to identify pathways and biological processes was performed on g:Profiler available at https://biit.cs.ut.ee/gprofiler/, with entities having g:SCS threshold < 0.05 considered significant [74]. Visualization was done using ggplot2 [75], except for Figure 6 generated using EnrichmentMap v.3.5.0 plugin [76] in Cytoscape v 3.10.3 [77].

## Data Availability

The results published here are in whole or part based upon data generated by the TCGA Research Network: https://www.cancer.gov/tcga. Controlled-access germline variants of TCGA cohorts were downloaded from the Genomic Data Commons (GDC, https://gdc.cancer.gov/about-data/publications/PanCanAtlas-Germline-AWG) of the National Cancer Institute (NCI) per Huang *et al.* [30]. TCGA variant and meta-data are available through the GDC portal at https://portal.gdc.cancer.gov/. Ancestry categories were obtained from TCGAA [33] available at http://fcgportal.org/TCGAA/. We chose to not analyze samples labeled “Native American [NA]” out of respect for Indigenous sovereignty.

Genes with germline associations at the tissue-specific level were downloaded from the COSMIC v90 [78] census list’s ‘Germline’ column. Full gene lists (n = 2 to 11) are available in Supplementary Table 3.

## Code Availability

We used PLINK v1.90 [79] available at https://zzz.bwh.harvard.edu/plink/ to calculate IBS matrices used to generate MDS plots. DBSCAN clusters were generated using the “dbscan” package in R (https://cran.r-project.org/web/packages/dbscan/index.html) [80], and HClust and K-Means using “stats” (https://stat.ethz.ch/R-manual/R-devel/library/stats/html/00Index.html).

## Acknowledgements

No acknowledgements to include in this section.

## Author Contributions

Conceptual basis: HM. Methods and experimental design: HM, KB and LP. Experiment execution and data analysis: HM. Discussions: HM, KB, PE, QM, JCZ, and LP. First manuscript draft: HM and LP. Final manuscript: HM and LP with input from KB, PE, QM, and JCZ.

## Competing Interests

LP has received consulting fees and honoraria for advisory board participation from Pfizer, Astra Zeneca, Merck, Novartis, Bristol-Myers Squibb, GlaxoSmithKline, Genentech, Personalis, Daiichi, Natera, Exact Sciences and institutional research funding from Seagen, GlaxoSmithKline, AstraZeneca, Merck, Pfizer and Bristol Myers Squibb.

## Notes

### Summary of Updates

Expanded results with respect to the number of cancer types and the clinical and functional biological analyses.

