## Supplementary Figures for "Dynamic clustering of genomics cohorts beyond race, ethnicity—and ancestry"

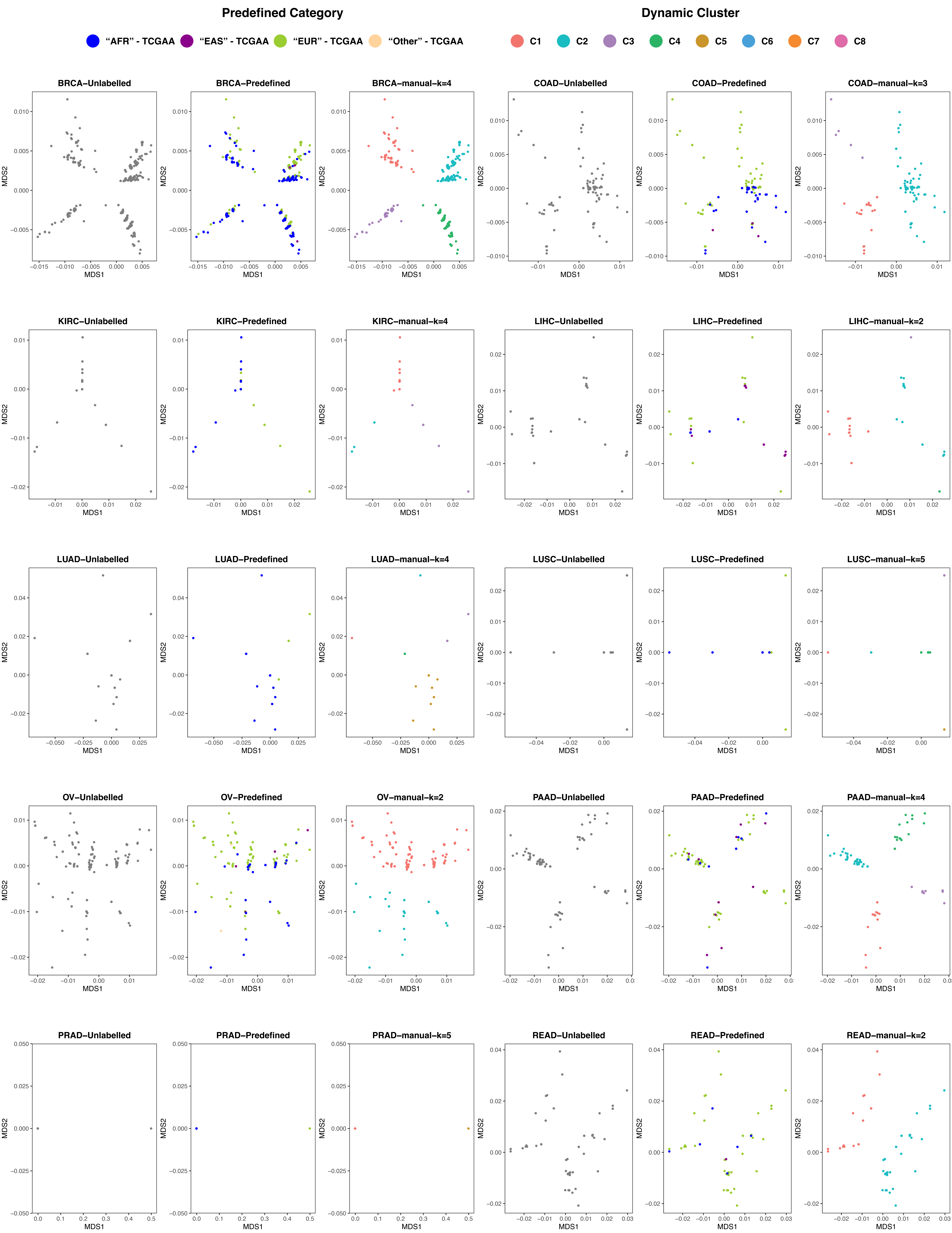


**Supplementary Figure 1.** HFI-based dynamic clustering across cancer types. Colors correspond to: no labels, predefined categories, and selected clusters after visual inspection.

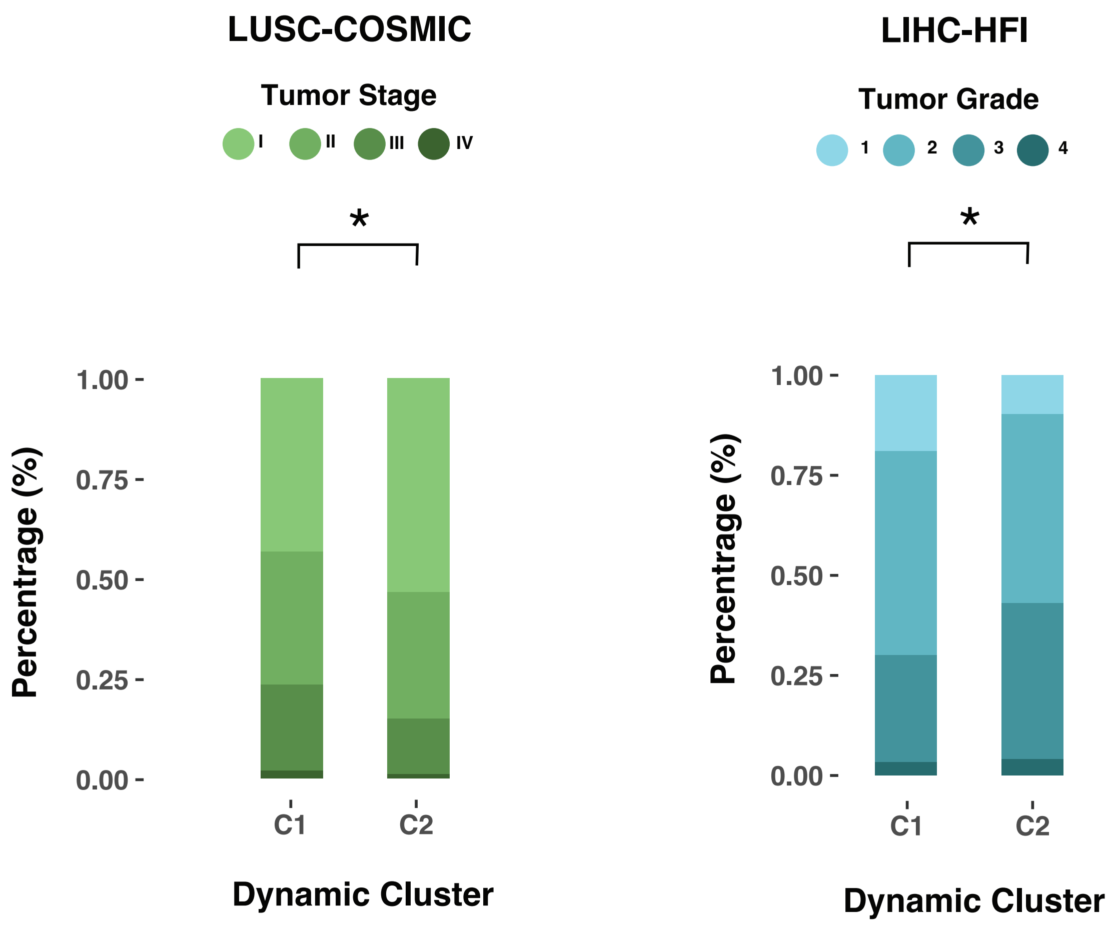


**Supplementary Figure 2.** Clinical associations with dynamic clusters in LUSC-COSMIC and LIHC-HFI settings.

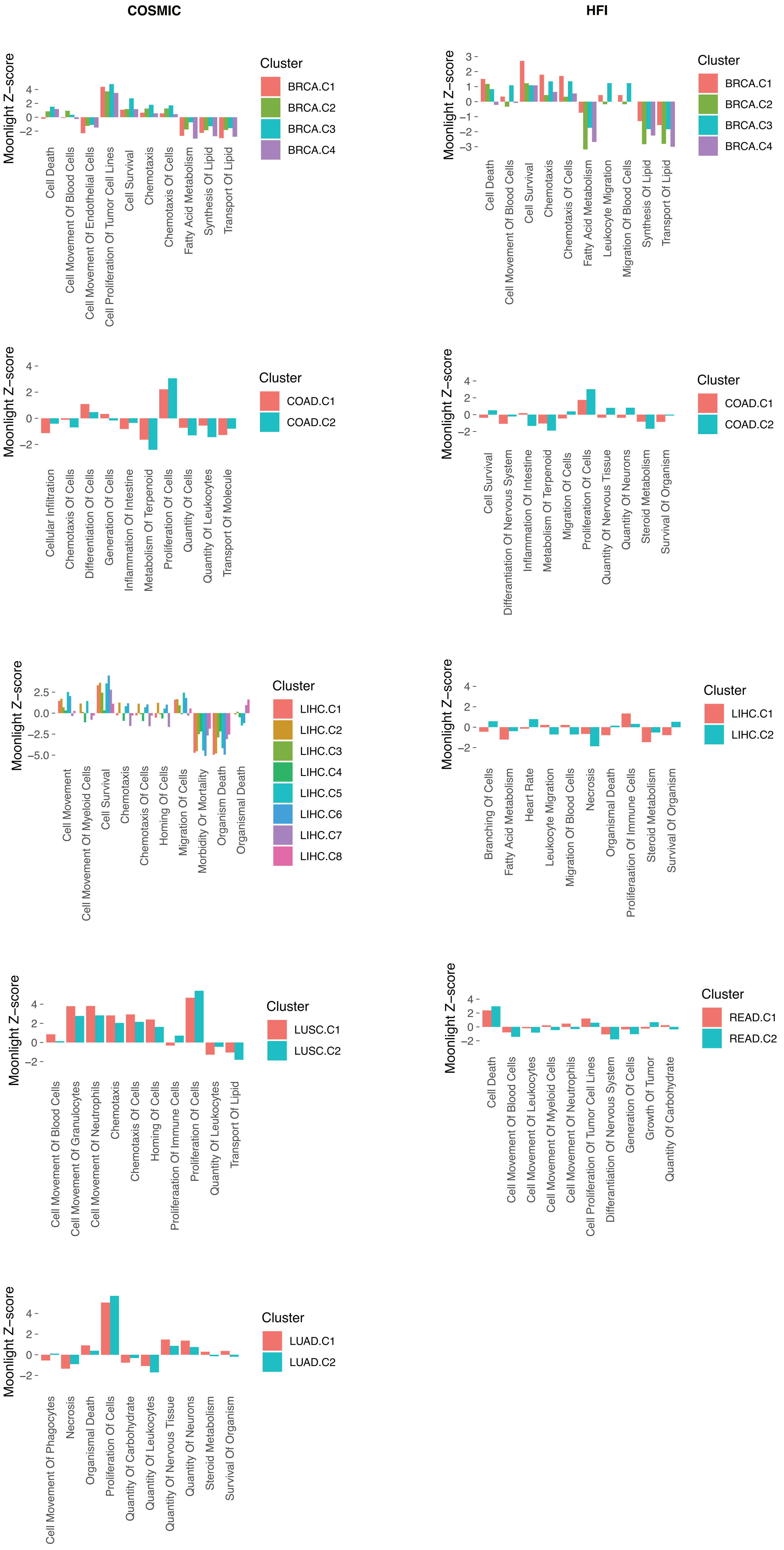


**Supplementary Figure 3.** Top Moonlight results of differentially expressed gene programs across cancer types and settings.

### LIHC-HFI-C1

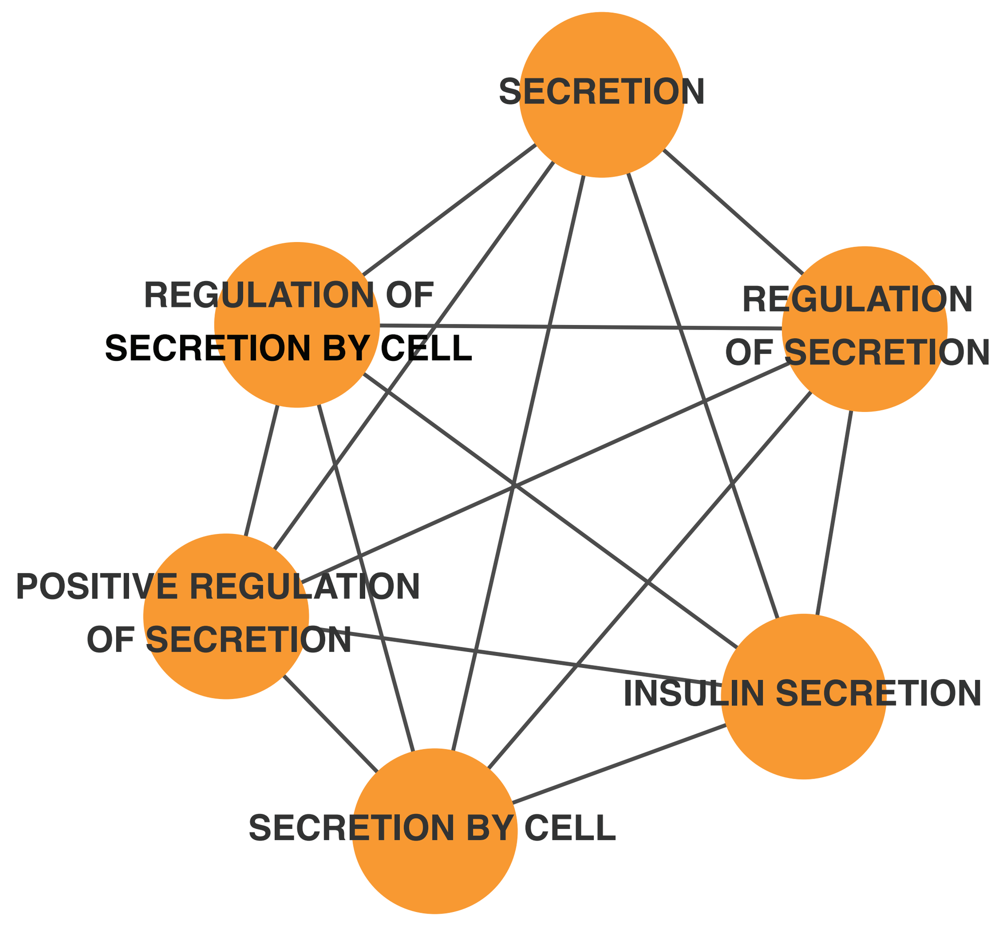


### LUSC-COSMIC-C2

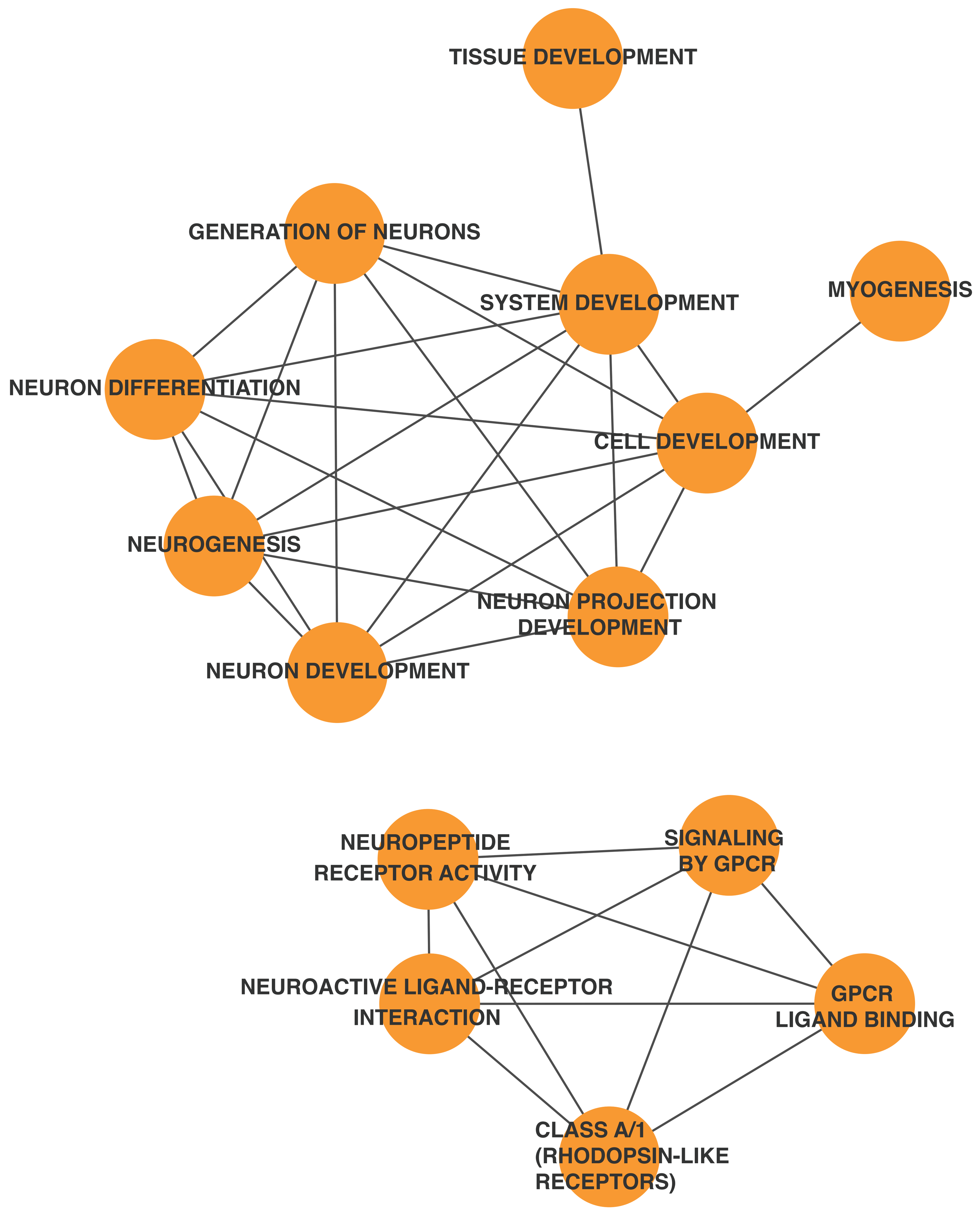


### READ-HFI-C2

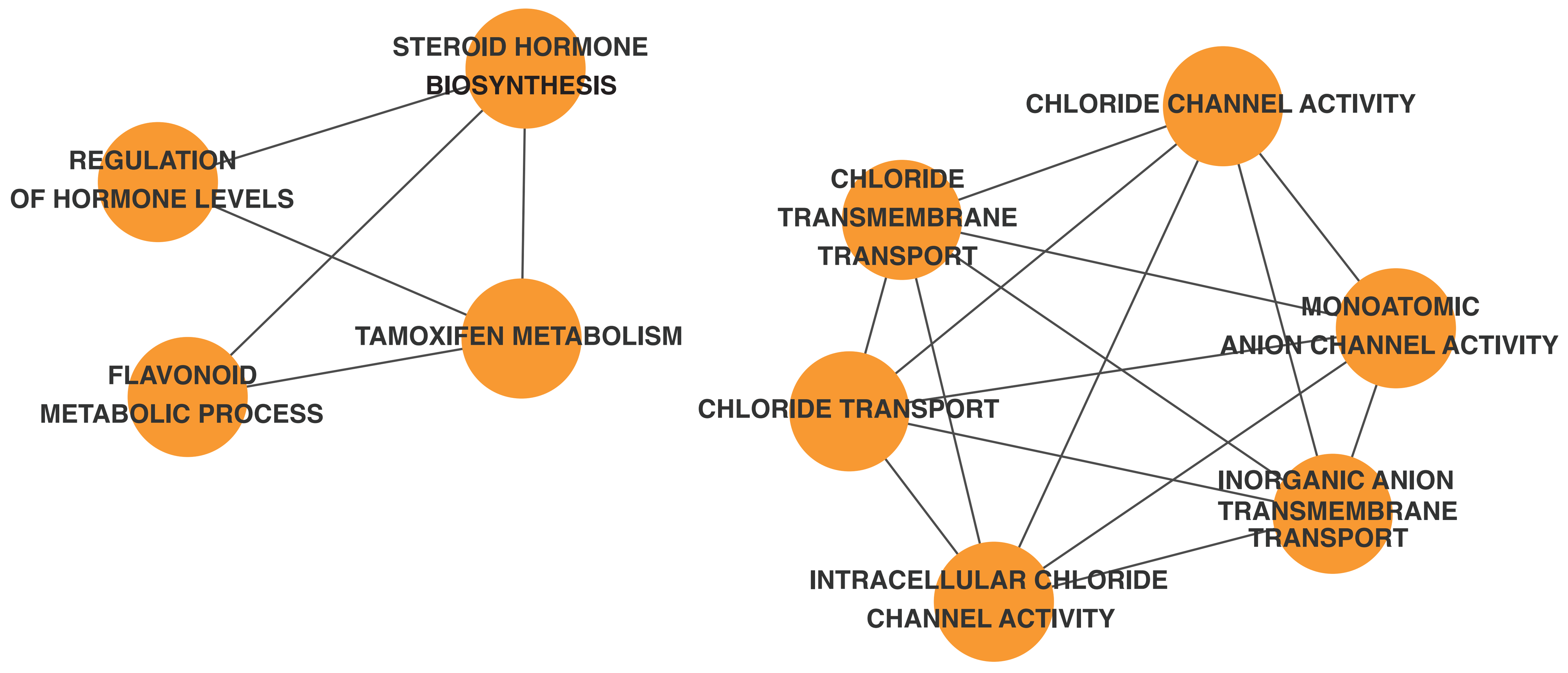


**Supplementary Figure 4.** Non-cancer signifiers significantly associated with single dynamic clusters in LIHC-HFI-C1, LUSC-COSMIC-C2, and READ-HFI-C2
